# Net radiation estimation using the Brunt equation for clear-sky emissivity and air and canopy temperatures for longwave radiation in well-watered crops

**DOI:** 10.64898/2026.03.31.715568

**Authors:** Thiago F. Duarte, Xuejun Dong, Daniel I. Leskovar, Uzair Ahmad, Noemi Tortorici, Tonny José Araújo da Silva, Edna Maria Bonfim da Silva

## Abstract

Net radiation (R_n_) can be estimated using models that apply the Brunt equation for the incoming longwave radiation and air temperature (T_air_) for the outgoing longwave radiation under reference conditions. This study aimed to estimate R_n_ using two previously regionally calibrated Brunt model, thereby eliminating the need site-specific calibration, and to assess whether T_air_ can be used as a substitute for canopy temperature (T_c_) under well-watered crop conditions. Measurements were conducted in sesame and cotton fields during the first year and in a cotton field during the second year. Canopy temperature was measured during the second year, and the calculations were performed at hourly and daily time scales. Regardless of the method used to estimate sky emissivity or whether T_c_ or T_air_ was used, errors were greater at hourly time scale. The overall RMSE, MAE, Bias and KGE values at the daily time scales were 11.88, 9.13, 2.53, and 0.91, in the first year, and 13.45, 10.56, 0.10 and 0.74, in the second year, respectively. When using both regionally calibrated Brunt model, R_n_ simulation performance was superior to that of the Allen/FAO method. The comparison between R_n_ estimated using T_air_ and T_c_, indicated statistical differences. Nevertheless, linear regression and error metrics showed that these differences were modest, especially at daily time scale. Thus, for practical purposes both regionally calibrated Brunt equations can be used to calculate clear-sky emissivity and improve R_n_ estimations, and T_air_ can be used as a substitute for T_c_ at the daily time scale under well-watered conditions.

## 1. Introduction

Net radiation (R_n_) results from the radiation balance at a specific site, where the downward radiation fluxes reaching the ground surface are balanced by the upward reflected shortwave radiation and emitted longwave radiation from the surface. It represents the amount of energy available to drive biophysical processes such as latent heat flux, sensible heat flux, soil heat flux. Consequently, R_n_ is generally required as an input variable for biophysical models based on the energy balance concept. For example, R_n_ is a required input variable for all the following models: the canopy stomatal conductance model proposed by Blonquist Jr. et al. (2009), the original crop water stress index (CWSI) model derived by Jackson et al. (1981), the Penman-Monteith equation (Allen et al. 1998), and the approach for modeling the soil temperature developed by Holmes et al. (2008).

Despite its importance, R_n_ is not always measured directly, mainly because it is expensive and difficult to measure accurately (Blonquist Jr. et al. 2010). In such cases, R_n_ must be estimated using theoretical or empirical models. The basic theoretical equation for estimating R_n_ consists of two components: net shortwave radiation (S_n_) and net longwave radiation (L_n_). The calculation of S_n_ is usually straightforward, as it depends on the incoming shortwave radiation and surface albedo – data that are commonly available. However, calculating the L_n_ component is more challenging because it depends on the atmospheric composition and temperature, which are needed to estimate the incoming longwave radiation, as well as on surface temperature, which is required to estimate emitted longwave radiation – variables that are not routinely available.

When modeling the incoming longwave radiation, sky emissivity (□_sky_) is typically estimated based on near-surface air temperature and vapor pressure, and sky temperature is replaced by the air temperature at the same reference height (Jensen and Allen 2016). Sky emissivity is calculated using clear-sky emissivity models (□_c_) adjusted by cloud fraction, since the presence of clouds significantly increases sky emissivity. Several models have been developed to calculate □_c_, as reviewed by Flerchinger et al. (2009), Formetta et al. (2016) and Li et al. (2017), and models’ performance is generally improved after proper calibration. However, Li et al. (2017), after working with fifteen models to estimate □_c_ in United States, concluded that although these models have different functional forms and coefficients, most of them exhibit similar accuracies after calibration and can be grouped into only two-families of models, the Brunt type (Brunt 1932) and the Carmona type (Carmona et al. 2014). Moreover, because the calibrated Brunt model has the simplest functional form and is among the most accurate, it can be recommended as a reference for calculating □_c_.

Similarly to sky temperature, surface temperature is not routinely measured, and its substitution is complex and should be applied with caution, as it depends on several factors such as ground cover type, soil type, soil water content, and others. Under reference conditions, as described by Jensen and Allen (2016), where the surface evaporates and transpires at rates corresponding to reference evapotranspiration, surface temperature (T_s_) is sufficiently close to air temperature (T_air_); therefore, T_s_ can be approximated by T_air_. For non-reference conditions, however, this approach is generally not recommended, particularly when the surface is dry and hot. Alternatively, surface temperature can be simulated using numeric methods, such as Newton’s iterative method (Bristow, 1987) or the iterative solution of aerodynamic fluxes (Jensen and Allen 2016). Nevertheless, these approaches significantly increase the complexity of R_n_ estimation, especially at shorter time-scales, as they require air-stability correction for calculating aerodynamic resistance as well as proper estimation of bulk surface resistance. In addition, using simulated T_s_ may introduce errors that could propagate uncertainty in the R_n_ estimation. Thus, in cases where a simple approach of approximating T_s_ by T_air_ is suitable, it should be preferred for R_n_ estimation.

In this sense, for example, An et al. (2017) reported that substituting soil temperature for air temperature in net radiation estimates over an experimental embankment did not significantly affect the results at half-hourly and hourly scales. Conversely, Linacre (1969) in his classical work, found that for monthly estimates of net radiation, approximating T_s_ by T_air_ led to significant errors across different soil types. More recently, Pieri (2010) developed a net radiation partition model applied to grapevines and reported that substituting leaf temperatures with air temperature measured near the canopy had only a minor impact on model performance.

In this study, net radiation was measured in irrigated cotton and sesame fields over two years: 2015 (sesame and cotton) and 2025 (cotton). In the second year, canopy temperature was also measured, beginning at the nearly full canopy stage. Under this condition, we assumed that the surface temperature could be represented by the canopy temperature (T_c_). We then tested whether surface temperature could be replaced by air temperature, as is commonly assumed in net radiation studies involving reference crops. Furthermore, the Brunt-type model calibrated by Formetta et al. (2016) and Li et al. (2017) for estimating □_c_ was also evaluated. These studies were selected for two reasons. First, Formetta et al. (2016) performed calibrations for several U.S. states, including Texas – where this study was conducted – and generated regionally specific calibrated parameters. Second, to test whether a more general or universal calibration equation would result in accurate longwave radiation estimation, the model of Li et al. (2017) was selected, as their equation was derived from grouped data from a large dataset collected across the United States. We hypothesized that the R_n_ estimation can be improved by using a broadly calibrated □_c_ equation, thereby eliminating the need for site-specific calibration. We also hypothesized that, under well-watered conditions in cotton, T_s_ can be approximated by T_air_ without loss of accuracy in R_n_ estimation.

## 2. Materials and methods

### 2.1. Locality and measurements

The study was carried out in a farm field at the Texas A&M AgriLife Research and Extension Center at Uvalde. The local soil is Fine-silty, mixed, active, hyperthermic Aridic Calciustoll (https://soilseries.sc.egov.usda.gov/OSD_Docs/U/UVALDE.html), and the climate is classified as Humid Subtropical climate (Cfa) according to the Köppen climate classification system.

The net radiation measurements were performed in 2015 and 2025. In 2015, the measurements were taken in a 4.9-ha cotton crop field from 7/15/2015 to 9/4/2015 and in a 4.9-ha sesame crop field from 9/4/2015 to 10/29/2015. The two crop fields were located about 80 meters from each other within a 20-ha field irrigated using a center pivot system. Cotton (variety PHY 499 WRF) was cultivated from 04/02/2015 to 09/23/2015, with 1.02 m row spacing and a stand of 100,000 pl ha^−1^, and during the net radiation measurements the leaf area index changed from 4.85 m^2^ m^−2^ to 3.49 m^2^ m^−2^. Sesame (variety S36 from Sesaco Co. San Antonio, TX, USA) was cultivated from 7/16/2015 to 11/23/2015, with 1.02 m row spacing and a stand of 450,000 pl ha^−1^ with the leaf area index changing from 4.86 to 1.49 m^2^ m^−2^ during the measurements (Figure 1). In 2025, the net radiation measurements were taken only in a cotton field, from the period of 6/19/2025 to 9/02/2025, with the leaf area varying from 1.5 to 3.8 m^2^ m^−2^ (Figure 1). Similarly to 2015, the cotton (varieties NG 4190 and ST 4990) was planted with 0.76 m row spacing and a stand of 129,000 pl ha^−1^. Leaf area index for the cotton crop in both years was measured using a LI-2200C Plant Canopy Analyzer (LI-COR, Inc. Lincoln, NE, USA) on relatively calm days without significant amounts of broken clouds in the sky. The LAI for the sesame crop was estimated using the field-measured data of oven-dried green leaf mass per unit land area multiplied by the specific leaf area.

**Figure 1.**
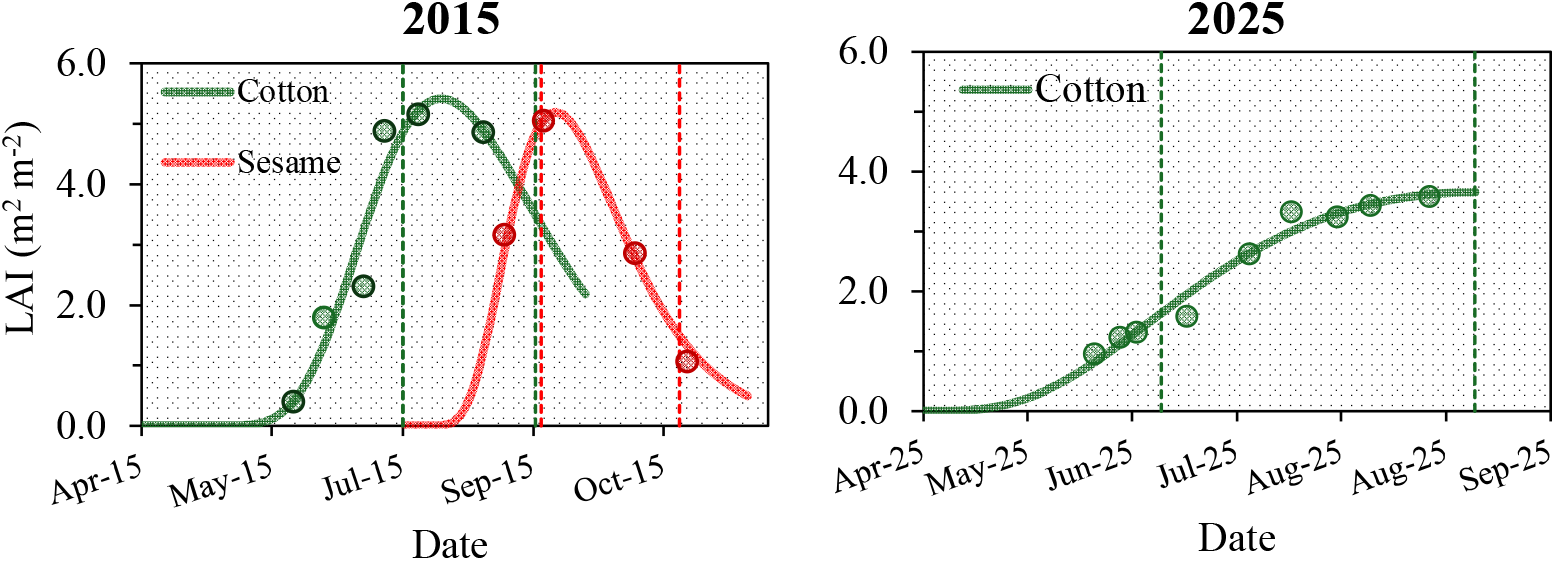
Leaf area index variations in cotton and sesame plots during the crop cycle in which the net radiation was measured in 2015 and 2025 at the Uvalde Research Center. The vertical dashed lines represent the period of net radiation measurement.

In both years, the crops were irrigated to meet crop water demand, calculated based on maximum evapotranspiration. In 2015, a center-pivot irrigation system was used, while a dripline system was used in 2025.

In both 2015 and 2025 the net radiation measurements were performed using an NR-Lite2 Net Radiometer sensor (Kipp & Zonen, Delft, The Netherlands) installed at a height of 3.0 m above the ground surface, with data collected at 1-minute intervals. In 2015, the data was recorded using a LOGBOX SD datalogger (Kipp & Zonen, Delft, The Netherlands), while in 2025 the measurement was made using a CR1000 datalogger (Campbell Scientific, Logan, Utah, USA). Additionally, solar radiation (W m^−2^), air temperature (°C) and relative humidity (%), were measured at 5-minute intervals at an automatic weather station located within 500-m from the experimental site. In 2025, canopy temperature was also measured using a set of infrared radiometers (SI-111-SS and SI-131-SS; Apogee Instruments, Inc., Logan, Utah, USA) at 15-minute intervals via a CR1000 datalogger. Measurements were obtained using two sensors installed approximately 50 cm apart and 15 cm above the top of the canopy. The sensors were tilted at approximately a 35° angle relative to the vertical axis, with one sensor directed toward the west side, and the other toward the east side on one representative cotton row (Figure S1). One advantage of tilting the sensors was to avoid soil surface exposure to the sensors’ field of view. The average of the two measurements was then calculated and used to estimate the longwave radiation emitted by the cotton canopy, as described in the next section.

### 2.2. Net radiation models

The basic equation to estimate R_n_ can be described as below:

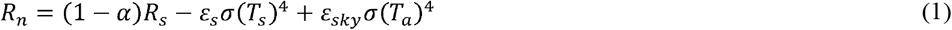

where R_n_ = net radiation (W m^−2^); α = surface albedo, defined as 0.21; R_s_ = solar radiation (W m^− 2^); □_s_ = surface emissivity; □_sky_ = sky emissivity; σ = Stefan-Boltzmann constant (5.670 × 10^−8^ W m^−2^ K^−4^); T_s_ = surface temperature (K); T_a_ = air temperature (K).

Equation 1 can be simplified to the following equations:

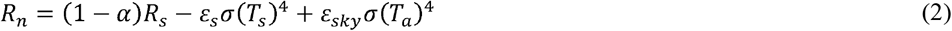

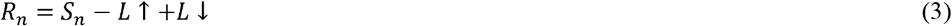

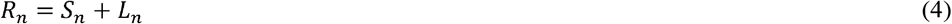

where S_n_ = net shortwave radiation (W m^−2^); L↑ = upward longwave radiation (W m^−2^); L↓= downward longwave radiation (W m^−2^); L_n_ = net longwave radiation (W m^−2^).

In this work, net radiation was calculated in hourly and daily time scales using different approaches described in detail in the following sections. The difference between the methods lies in how L_n_ was calculated, either by altering L↓ or by altering L↑. The method proposed and described by Allen et al. (1994) was considered the standard method. In 2015, as canopy temperature measurements were not available, only different L↓ models were tested. In 2025, the L↑ component was estimated based on measured canopy temperatures, and L↓ was also tested using different models.

#### 2.2.1. The FAO-Allen et al. (1994) model

In the Allen et al. (1994) model, net radiation is estimated in the daily time scale using equation 5 and its components.

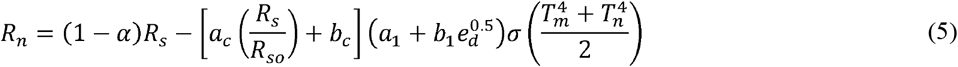

where R_n_ = net radiation (W m^−2^); α = surface albedo, defined as 0.21; R_s_ = solar radiation (W m^− 2^); R_so_ = clear-sky solar radiation (W m^−2^); e_d_ = vapor pressure at dew point (kPa); T_m_ = maximum air temperature (K); T_n_: minimum air temperature (K); a_c_ and b_c_ are the cloud factors, = 1.35 and –0.35, respectively (-); a_1_ and b_1_ = area emissivity factors, defined as 0.35 and – 0.14; σ = Stefan-Boltzmann constant (5.670 × 10^−8^ W m^−2^ K^−4^).

The variables e_d_ and R_so_ can be calculated using equations 6 and 7, as follows:

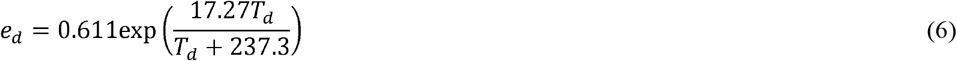

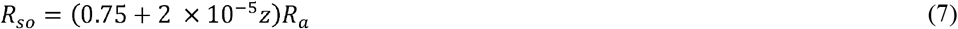

where T_d_ = dewpoint temperature (°C); z = is elevation above the sea level (m); R_a_ = extraterrestrial solar radiation (W m^−2^). The R_a_ in turn is calculated using equations 8, 9 and 10.

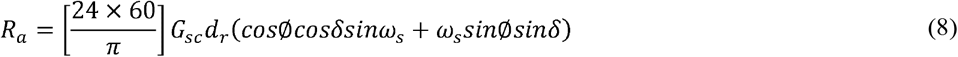

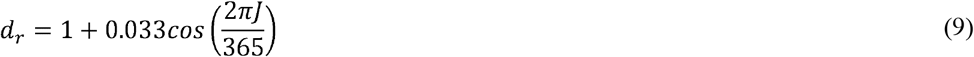

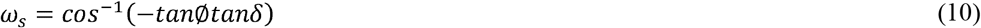

in which G_sc_ is the solar constant (1367 W m^−2^); d_r_ is the relative distance between the Earth and the sun (1.496 × 10^11^m); J is the Julian day of year; ω_s_ is the sunset time angle (rad); □ and δ are the latitude (rad) and solar declination (rad).

The previous equations can be adjusted to calculate the R_n_ in shorter timescale (An et al. 2017, Allen et al. 1994):

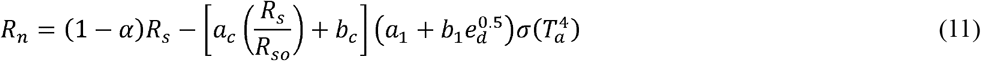

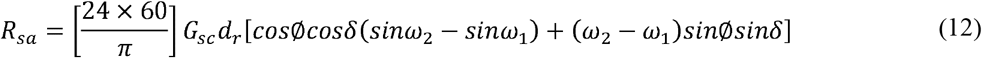

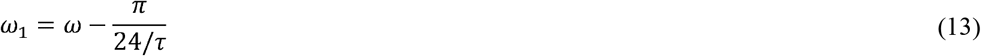

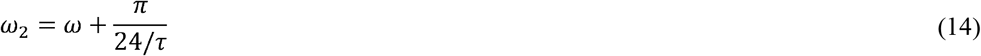

where T_a_ = mean air temperature (K); ω_1_ and ω_2_ are the solar time angles (rad) at the beginning and end of the considered period; ω is the solar time angle (rad) at the center of the period; and τ is the length of the period (h).

In Equations 5 and 11, the term 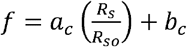 represents the cloudiness function. The approach of Snyder (2008) was applied in this study with the restriction of 0.3 < Rs /R_so_ ≤ 1.0 and Rs/Rso = 0 whenever β <17.5°. Furthermore, an initial *f* value of 0.6 was used, and for each sequential data, whenever β < 17.5°, the *f* value was set equal to the previous *f* value, and β represents the solar altitude:

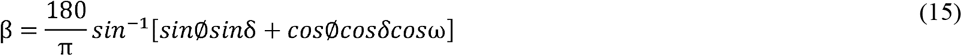

#### 2.2.2. Testing different sky emissivity models

The main challenge for the calculation of downward longwave radiation is the determination of the sky emissivity. In this work we tested the Brunt model for calculating clear-sky emissivity as calibrated by Formetta et al. (2016) and Li et al. (2017), as presented in Table S1. As pointed out before, these references were selected because they provide two options for □_sky_ calculation: one that is more regionally specific and another that is more universal, respectively.

Since the Brunt model is only valid for clear sky conditions, the correct estimation of downward longwave radiation for all-sky conditions requires considering the contribution of clouds. For this purpose, the Crawford and Duchon (1999) model were tested in this work.

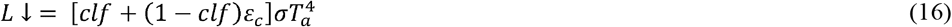

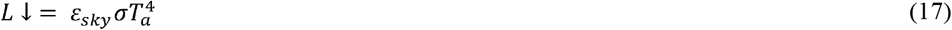

where clf is the cloud fraction term.

The clf was calculated as clf = 1 – s, where *s* represents the ratio between solar radiation measured at the weather station (W m^−2^) to the clear-sky radiation. The clear-sky radiation component can be calculated by several models, such as Jensen et al. (1990), Crawford and Duchon (1999), and Li et al. (2017). In this work the same approach used in equations 5 and 11, was used, that is, s = Rs/Rso.

#### 2.2.3. Estimating the upward longwave radiation

This approach differs from the previous two in that it uses the canopy temperature measurements to calculate upward longwave radiation and it was tested only with the 2025 data. Thus, the downward longwave radiation was calculated using the same equations described in the previous section, and the L↑ component was calculated as presented in Equations 2 and 3. The parameter □_s_ was set to 0.96, a representative value for the cotton crop (Campbell and Norman, 1998).

### 2.3. Statistical analysis

The net radiation methods described in the previous section were evaluated in hourly and daily. The measured and simulated data were compared using the Root Mean Square Error (RMSE), Mean Absolute Error (MAE) Bias and the Kling-Gupta Efficiency (KGE), as described below:

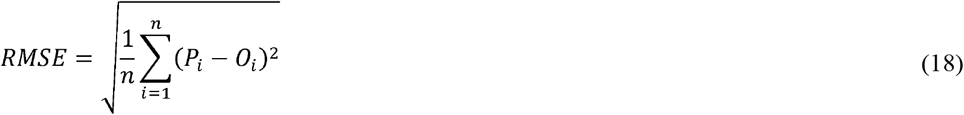

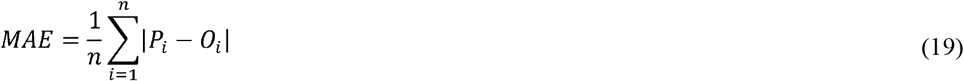

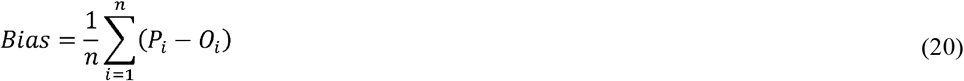

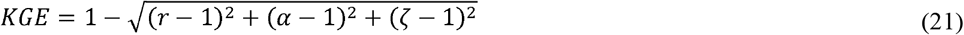

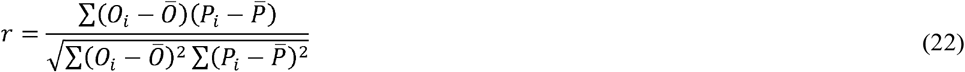

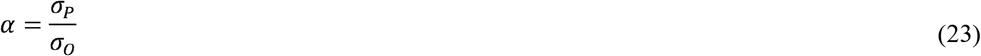

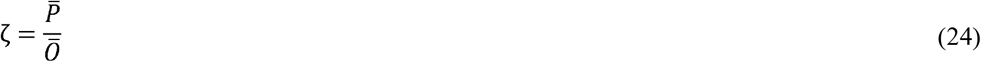

where: O_*i*_ is the observed value for the *i*-th data point; P_*i*_ is the predicted value for the *i*-th data point; ō is the mean of the observed values; 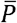 is the mean of predicted values; n is the total number of observations; σ_o_ is the standard deviation of observations, σ_P_ is standard deviation of predictions.

Additionally, the Wilcoxon signed-rank test and Bland–Altman analysis were used to compare R_n_ estimates derived from T_air_ and T_c_. The Wilcoxon test was applied to assess paired differences between methods using the Minitab 22, while agreement and systematic bias were evaluated using the Bland–Altman approach, calculated as described by Yellareddygari and Gudmestad (2017). The Bland-Altman analysis is a graphical method in which the differences between the R_n_ methods (Y-axis) and the averages of the methods (X-axis) are plotted on the same graph. The mean difference between the methods (i.e., Bias), and the limits of agreement, calculated as bias ± 1.96 × SD (SD – standard deviation), are also plotted. This method was first used in medicine (Bland and Altman, 1986), but more recently in crop and soil sciences studies (Yellareddygari and Gudmestad, 2017; Lipiec et al. 2021; Usowicz, et al. 2020).

## 3. Results and Discussion

The results of net radiation estimation using different approaches for 2015 are presented in Figures 2 and 3, with the corresponding statistical performance shown in Figure 4. It can be observed that the performance of the methods depends on the timescale, with more accurate values obtained on a daily scale than on an hourly scale. In general, considering all methods, the RMSE, MAE, Bias and KGE values were 38.95, 28.34, −5.49, 0.90 for hourly time scales, and 11.88, 9.13, 2.53, and 0.91 for daily time scales, respectively. Despite this, the R_n_ estimates using the approach of Formetta et al. (2016) for □_c_ were superior to those obtained with the other methods, with overall error across both crops of 37.74 (RMSE), 26.60 (MAE), −7.92 (Bias), and 0.94 (KGE).

**Figure 2.**
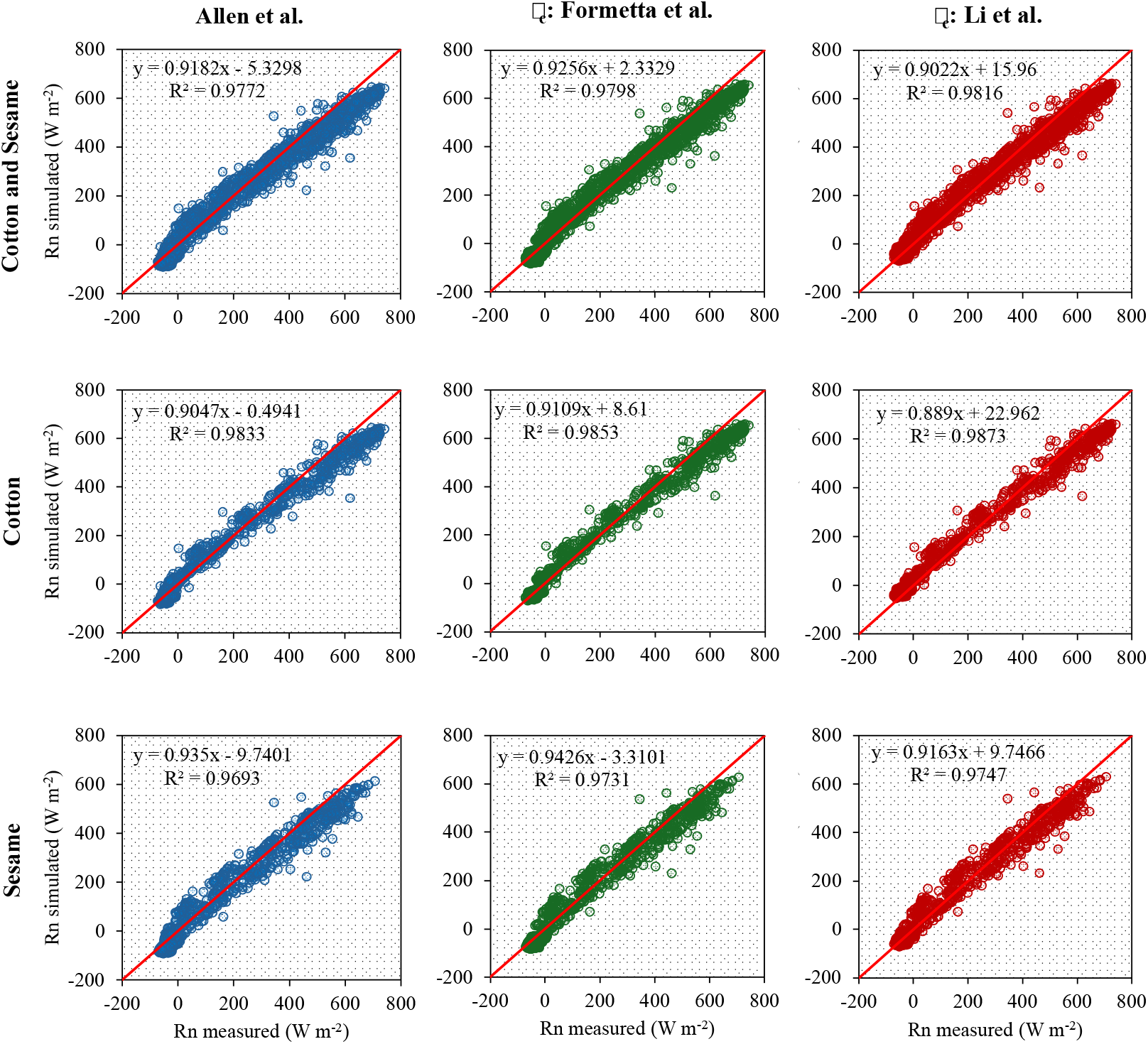
Relationship between hourly simulated and measured net radiation (W m^−2^) in cotton and sesame fields at Uvalde in 2015, using different methodologies. □_c_: clear-sky emissivity model.

**Figure 3.**
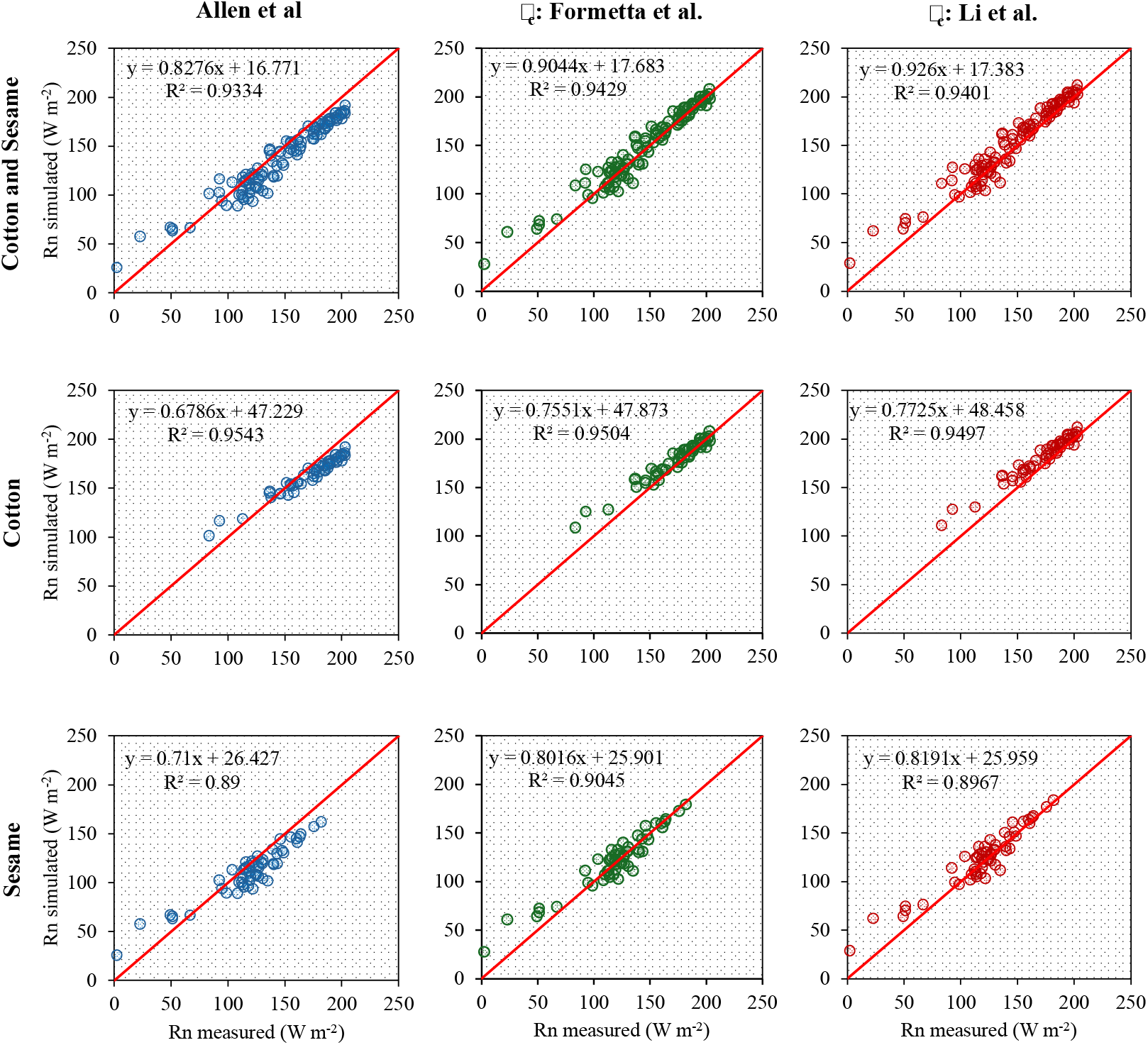
Relationship between daily simulated and measured net radiation (W m^−2^) in cotton and sesame fields at Uvalde in 2015, using different methodologies. □_c_: clear-sky emissivity model.

**Figure 4.**
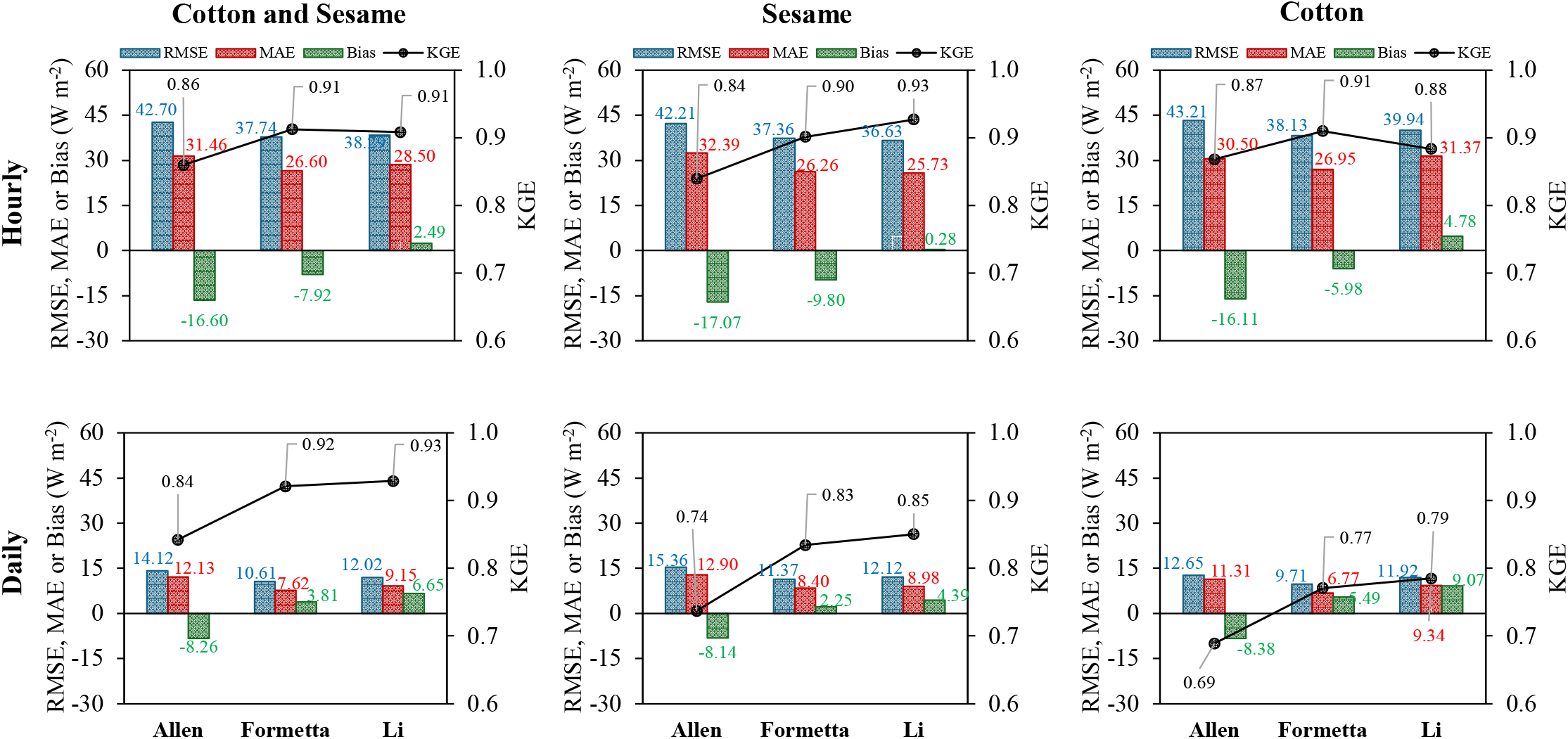
Statistical performance of net radiation calculation methods at hourly and daily time scales for the year 2015. Colored values represent the respective statistical metrics.

For the year 2025, the errors were slightly higher than those observed in 2015, particularly at the hourly time scale, as shown in Figures 5 to 7. The general errors were 55.94 (RMSE), 40.10 (MAE), −14 (Bias) and 0.81 (KGE) for hourly time scales, and 13.45, 10.46, 0.10, and 0.74 for daily time scales, respectively. Nevertheless, both methods used to estimate □_c_ outperformed the Allen method with its original calibrated parameters. The differences between the Formetta and Li methods were not very pronounced when using either T_air_ or T_c._ However, the Li equation consistently produced lower bias, regardless of the type of temperature that was used (Figure 7).

**Figure 5.**
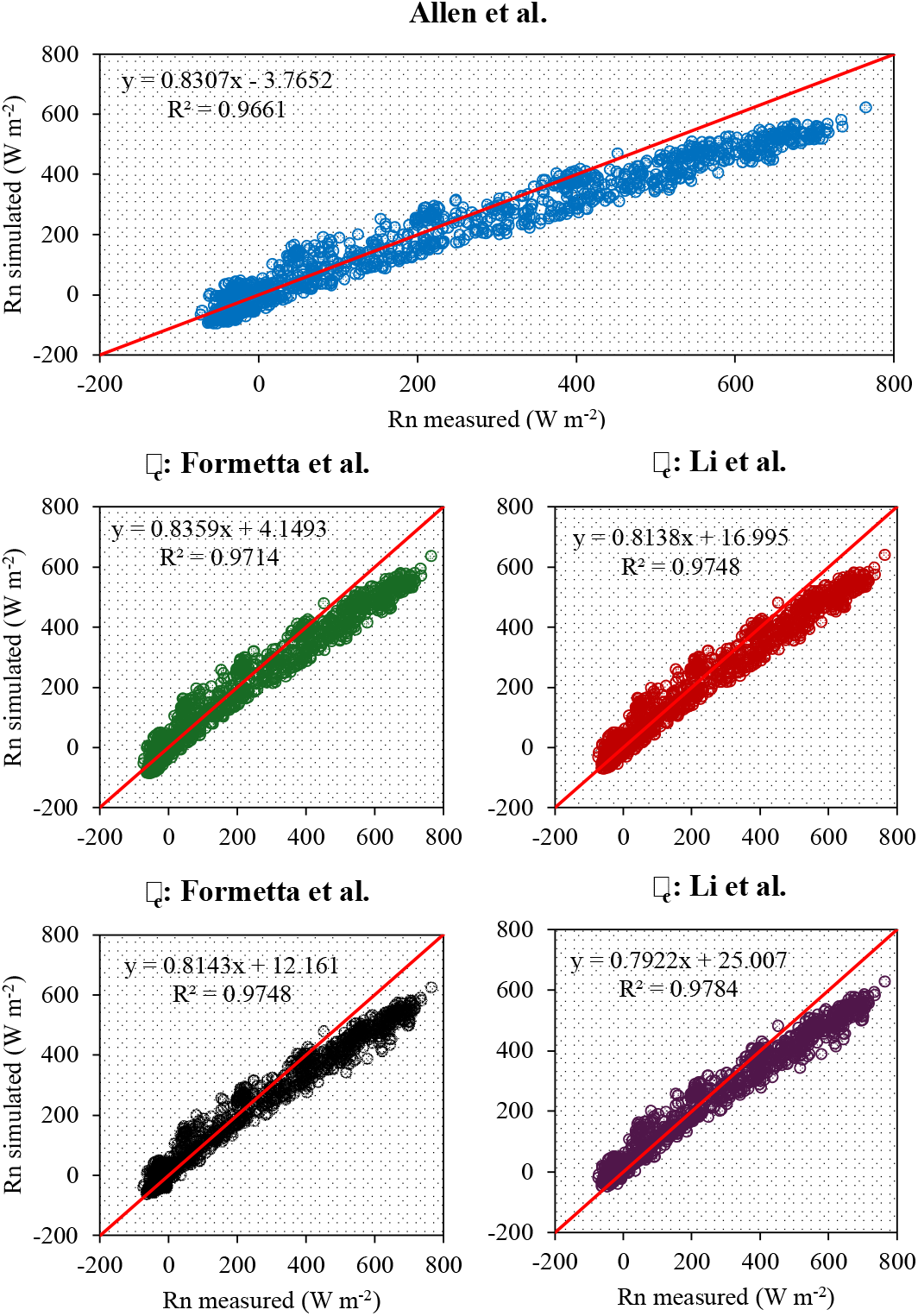
Relationship between hourly estimated and measured net radiation (W m^−2^) in cotton field at Uvalde in 2025, using different methodologies. □_c_: clear-sky emissivity model. T_s_: surface temperature (°C); T_air_: air temperature (°C); T_c_: canopy temperature (°C).

**Figure 6.**
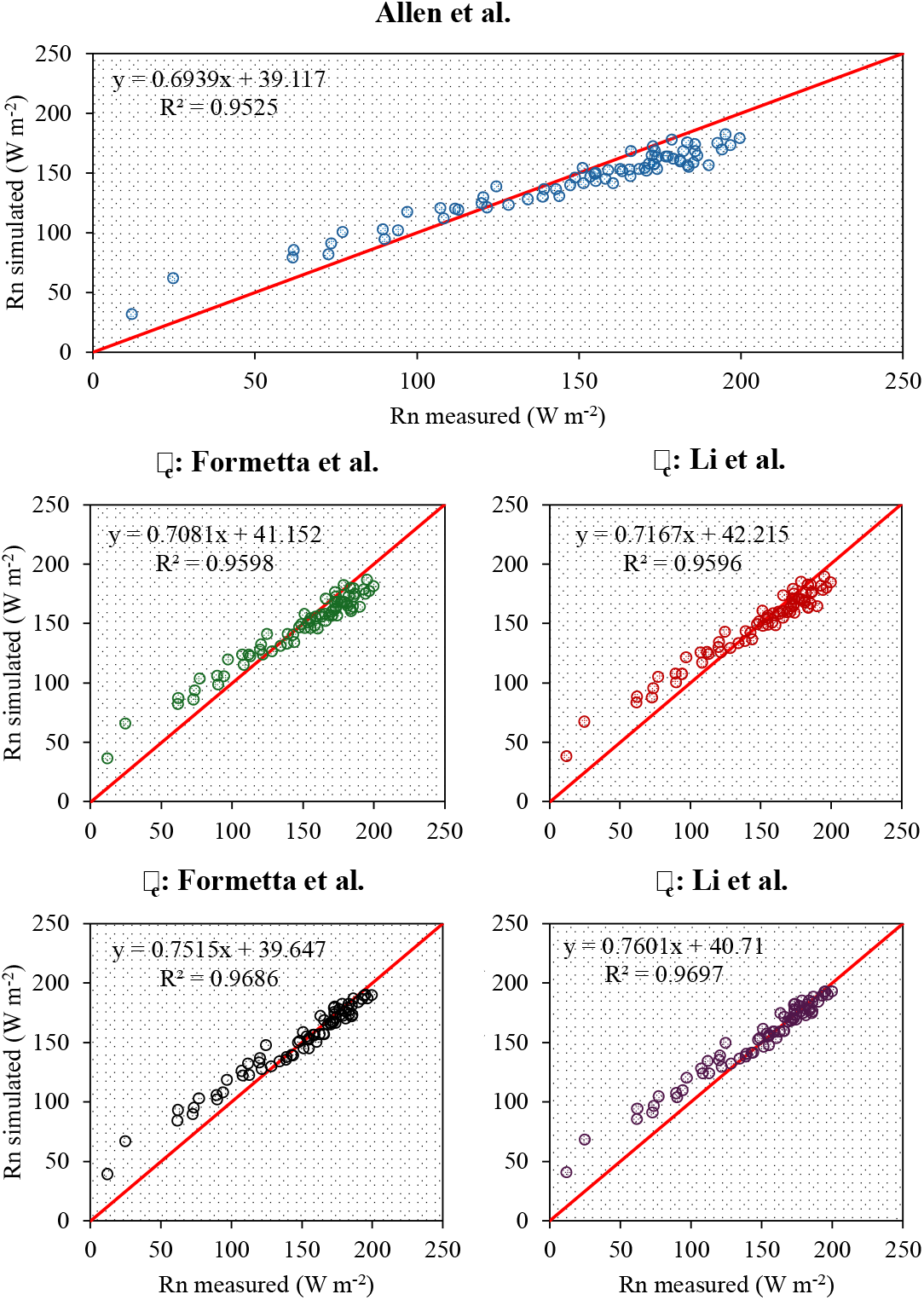
Relationship between daily estimated and measured net radiation (W m□^2^) in a cotton field during 2025, using different methodologies. □c: clear-sky emissivity model; Ts: surface temperature (°C); T_air_: air temperature (°C); Tc: canopy temperature (°C).

**Figure 7.**
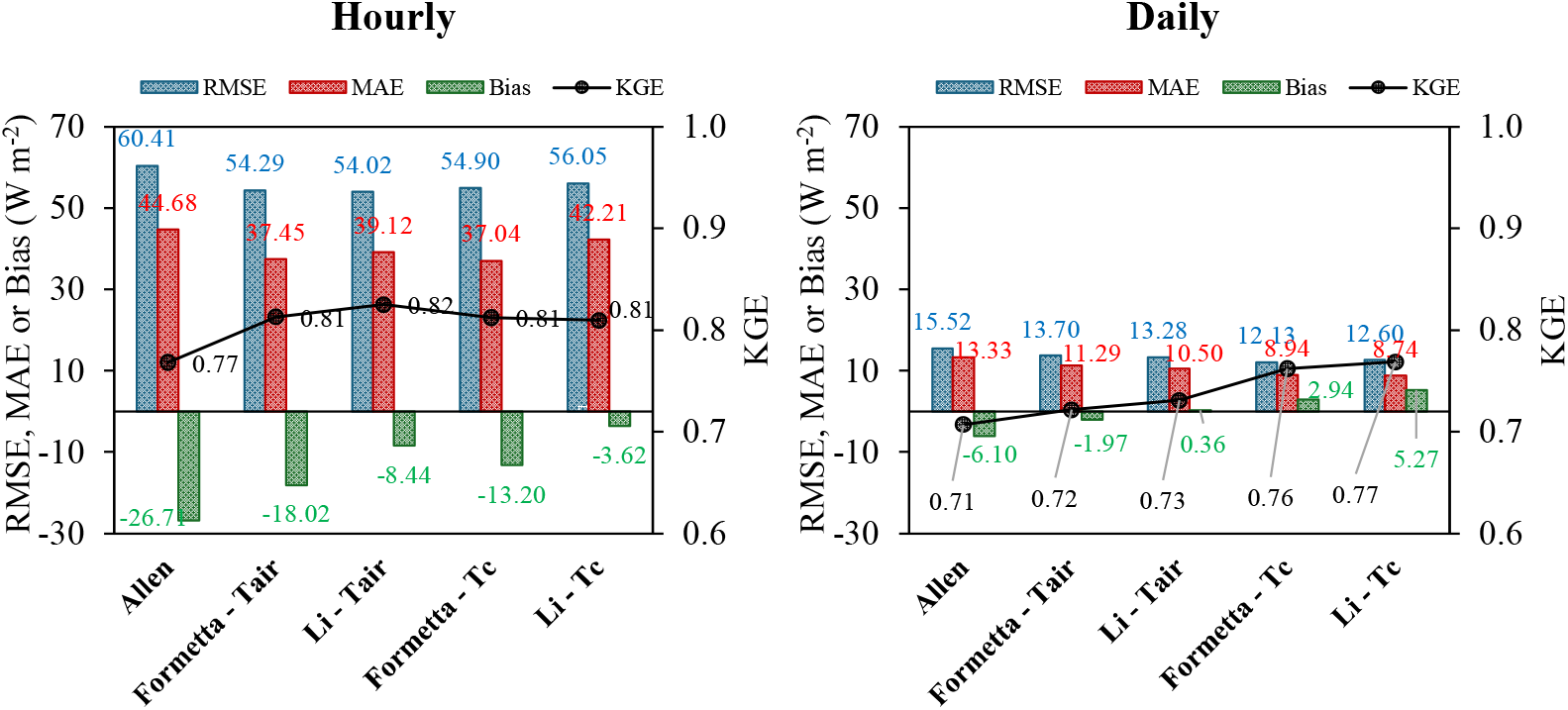
Performance of net radiation estimation methods across various time scales for the year 2025. Colored values represent the respective statistical metrics.

The higher errors in simulated R_n_ at shorter time scales (Figures 4 and 7) can be partially attributed to the assumption of constant albedo throughout the day. In turfgrass, for example, Blonquist Jr et al. (2010) found that albedo can vary from 0.21 to 0.30, while in cotton, data presented by Carvalho et al. (2022) showed that it can vary by approximately 0.08 units, changing from 0.22 to 0.30 at the flowering stage. However, Blonquist Jr et al. (2010) point out that the most critical errors in estimating R_n_ at short time scales are related to net longwave radiation. In this context, part of the error source can be attributed to the assumption that the cloudiness function remains constant during the night, which can lead to incorrect estimates for those periods. In addition, estimating atmospheric emissivity based on e_a_ and assuming that T_air_ is representative of atmospheric temperatures may not be suitable in some situations, and consequently may not accurately capture longwave radiation dynamics.

The performance of R_n_ estimation could likely have been improved with local, site-specific calibration, as demonstrated in other studies (Kjaersgaard et al. 2007; Myeni et al. 2020). However, one of the aims of this work was to evaluate whether previously calibrated coefficient could yield acceptable results, thereby eliminating the need for highly site-specific calibration. A similar attempt was made by Myeni et al. (2020) for southern African conditions, in which the authors tested whether R_n_ estimation could be improved using the original equations of Idso and Jackson (1969) or Brutsaert (1975) to estimate clear-sky emissivity in combination with the cloudiness factor model of Crawford and Duchon (1999). Overall, the authors found that the evaluated method outperformed the Allen model in its original form. In this work, we used the Brunt model calibrated by different authors, and although the parameter values differed, their performance was similar and superior to that of the original Allen equation. Thus, both approaches can be used to improve the estimation of clear-sky emissivity and, consequently, net radiation.

The higher errors observed in 2025 may be attributed to the lower LAI compared with 2015 (Figure 1), which resulted in reduced crop ground cover. Consequently, the fraction of exposed soil may have altered the amount of reflected shortwave radiation, thereby affecting net shortwave and, in turn, R_n_. A second source of error may also be attributed to the outward longwave radiation originating from the partially exposed soil surface, which exceeded the contribution accounted for in this study, since soil surface temperature normally differs from canopy temperature, and consequently alters the emitted longwave radiance.

Bsaibes et al. (2009), who analyzed the relationship between LAI and albedo for different crops, observed that, for low to moderate LAI values (LAI < 3), the soil background exerted a stronger influence on albedo values, whereas for larger LAI values, albedo tended toward an asymptotic value that may depend more on canopy characteristics such as architecture, like chlorophyll content, among others. Soil albedo is usually lower than the crop albedo, especially for dark and moist soils (He et al. 2019), with general values ranging from 0.08 (wet, dark soil) to 0.18 (soil, dry light), as pointed out by Campbel and Norman (1998). A higher soil contribution would result in lower shortwave reflectance and consequently, higher R_n_ values. In this study, we assumed a fixed albedo value of 0.21 for both years, which may have led to an underestimation of the actual R_n_ values.

To assess whether canopy temperature (T_c_) can be approximated by air temperature (T_air_) for R_n_ estimation, the R_n_ values calculated using both temperatures were compared using the Wilcoxon signed-rank test, as shown in Table 1. The results indicate that, for both time scales, the median difference between R_n_ estimated using T_air_ and T_c_ was significantly different from zero. Therefore, strictly speaking, for cotton crops, R_n_ cannot be estimated by substituting T_c_ with T_air_, even under well-watered conditions. However, overall, the error metrics were very similar between the two methods (Figure 7), with notable differences observed only in the Bias for hourly scale, ranging from −18.02 (T_air_) to −13.20 (T_c_) for Formetta, and from −8.44 (T_air_) to −3.62 (T_c_) for Li.

**Table 1.** Results of the Wilcoxon signed-rank test for comparing R_n_ estimates obtained from T_air_ and T_c_.

| Variable | n | Median | Wilcoxon statistic | p |
| --- | --- | --- | --- | --- |
| Hourly ($T_{air}$ vs $T_c$) | 1727 | -5.05 | 343890 | <.001 |
| Daily ($T_{air}$ vs $T_c$) | 72 | -4.52 | 139 | <.001 |

Nevertheless, linear regression analysis between R_n_ simulated using T_air_ and T_c_ showed that the regression line was close to the 1:1 line (Figure 8) for both hourly and daily time scales. For the hourly scale, the slope was close to 1, although the intercept was significantly different from zero, indicating a small systematic bias but, an overall good agreement (Table 2). For the daily scale, the intercept did not differ significantly from zero, and the slope was close to 1, demonstrating that differences between R_n_ estimated from T_air_ and T_c_ were negligible.

**Figure 8.**
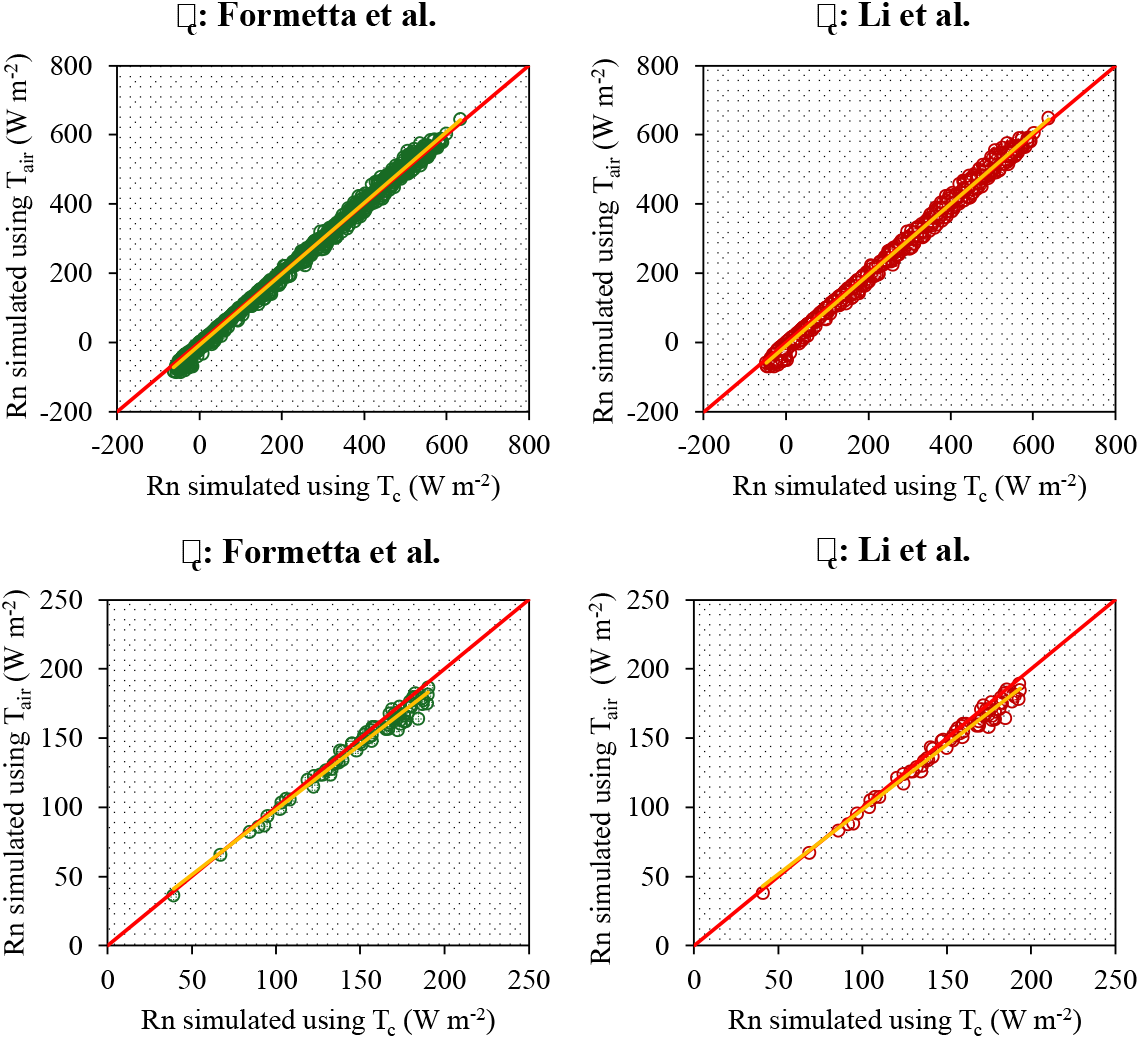
Linear regression comparing R_n_ estimates obtained from T_air_ and T_c_ with different models used to calculate □c. The red line indicates the 1:1 line, and the yellow line represents the regression line.

**Table 2.** Results of linear regression analysis comparing R_n_ estimates derived from T_air_ and T_c_ across different time scales and using various methods.

| Variable | Model | Intercept |  |  | Slope |  |  |
| --- | --- | --- | --- | --- | --- | --- | --- |
|  |  | Value | Lower<br>95% | Upper<br>95% | Value | Lower<br>95% | Upper<br>95% |
| Hourly ($T_{air}$ vs $T_c$) | Formetta | -8.427*** | -8.982 | -7.873 | 1.027*** | 1.025 | 1.029 |
|  | Li | -8.794*** | -8.221 | -9.366 | 1.028*** | 1.025 | 1.030 |
| Daily ($T_{air}$ vs $T_c$) | Formetta | 4.636<sup>ns</sup> | -0.381 | 9.653 | 0.937*** | 0.904 | 0.969 |
|  | Li | 4.566<sup>ns</sup> | -0.486 | 9.617 | 0.938*** | 0.906 | 0.970 |

\*\*\* Significant at $p < 0.001$ ; ns: not significant ( $p > 0.05$ ).

The Bland–Altman plot (Figure 9) revealed greater variability at the hourly scale compared to the daily scale. Although most data points were within the limits of agreement (95% of the differences), some estimates fell outside these limits, particularly for extreme R_n_ values. Conversely, at the daily scale, the errors between the methods appeared to be more uniformly distributed, with noticeable deviations occurring only at higher R_n_ values. It is interesting to note that the Bias and the limits of agreement are the same for both the Formetta and Li methods. This occurs because, in both cases, they use the same data to calculate R_n_ (T_air_ and T_c_); therefore, the difference in R_n_ calculated from T_air_ and T_c_ is expected to be the same. The bias at the hourly and daily scale were −4.82 W m^−2^ and −4.91 W m^−2^, respectively, indicating that estimating R_n_ using T_air_ results in an average difference of approximately – 5.0 W m^−2^ under well-watered conditions. Thus, for practical purposes—and considering the modest improvement obtained by using T_c_ instead of T_air_ (Figure 7)—the assumption of similarity may be particularly reasonable at the daily time scale, when the differences between T_c_ and T_air_ are also expected to be minimal (Figure 10).

**Figure 9.**
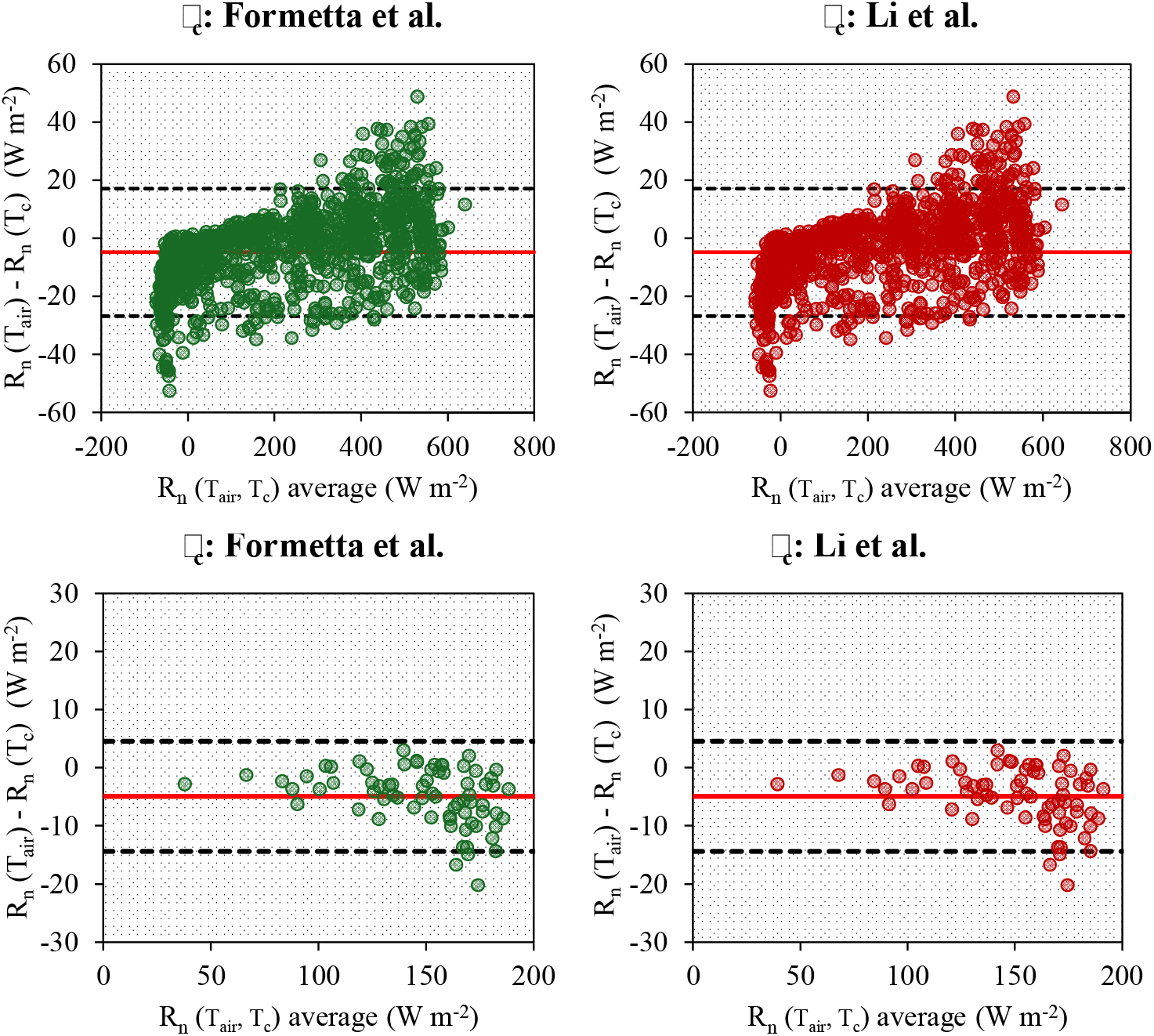
Bland–Altman plot comparing R_n_ estimates using T_air_ and T_c_, as well as different models for calculating □_c_. The upper and lower dashed lines indicate the limits of agreement, and the red line represents the Bias.

**Figure 10.**
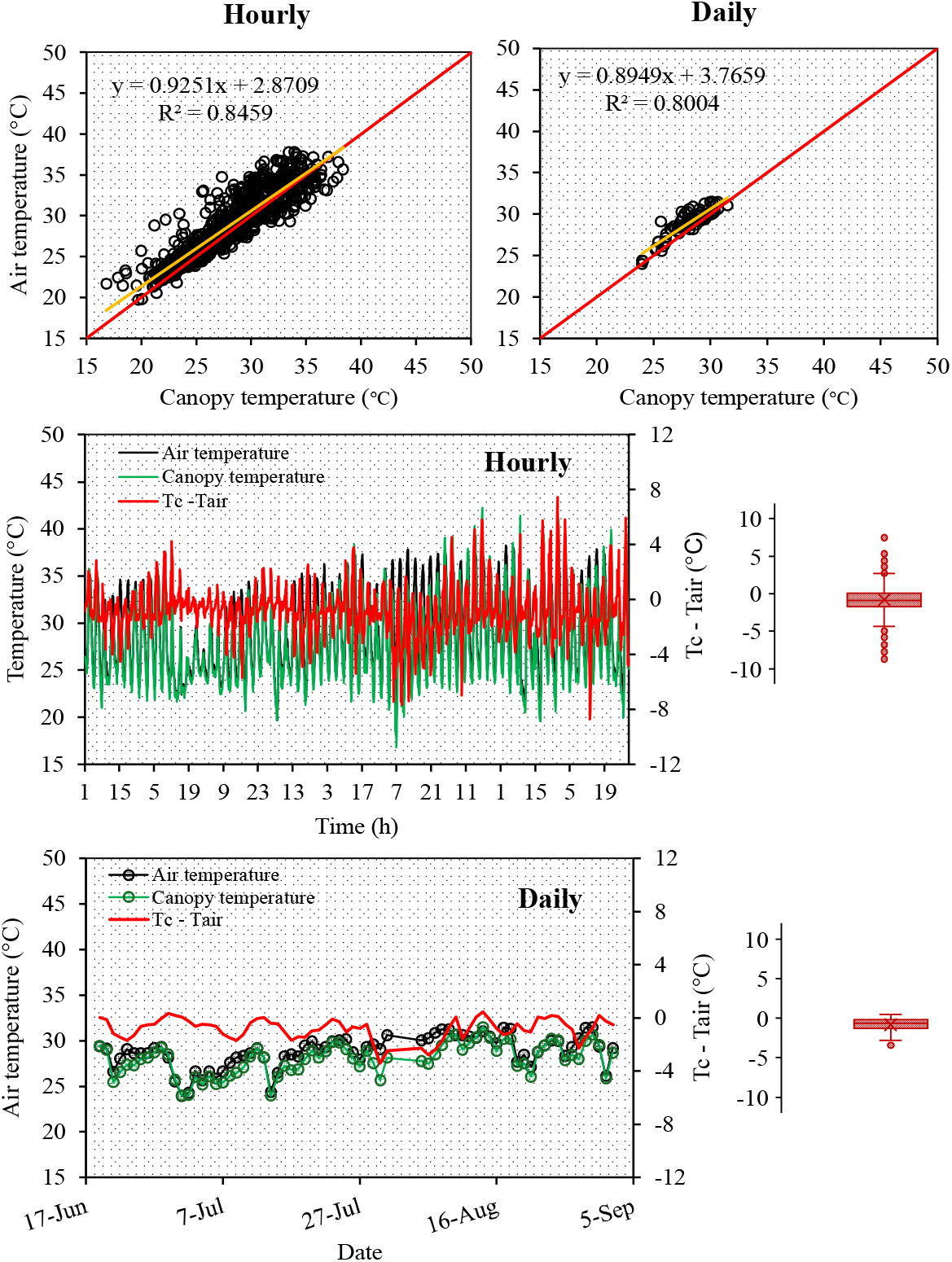
Relationship between air temperature (T_air_, °C) and canopy temperature (T_c_, °C) at hourly and daily time scales in a cotton field at Uvalde during 2025. The boxplot on the right shows the distribution of errors at each time scale.

The differences between T_air_ and T_c_ were largest at the hourly scale (Figure 10), as expected, with an average difference (T_c_ – T_air_) of –0.82 °C and a peak difference of –8.7 °C, indicating that T_air_ was generally slightly higher than T_c_. Nevertheless, at the daily scale, the differences were less pronounced, with a peak difference of –3.4 °C, which reinforced the use of T_air_ to estimate emitted longwave radiation at R_n_ estimation.

Although comparable studies in literature are scarce, similar results were reported by Carrasco and Ortega-Farías (2008), who evaluated two models to estimate R_n_ in vineyards (Vitis vinifera cv. Cabernet Sauvignon), considering either T_air_ as a substitute for T_c_ or the actual T_c_. Their results showed that the model using T_air_ was able to estimate net radiation with a good degree of precision, although the temperature gradient between the canopy and the air ranged from –3 to 4 °C, indicating that the incorporation of canopy temperature did not substantially improve R_n_ estimation, provided that soil water content were maintained at an optimum level to minimize the temperature differences.

This study focused exclusively on the substitution of T_air_ for T_c_ under well-watered conditions. However, no threshold was identified to define the temperature difference beyond which T_air_ and T_c_ can no longer be used interchangeably. This aspect requires further investigation, since R_n_ is also required in energy balance studies under non-irrigated or water-stressed crop conditions. The acceptable temperature difference threshold likely depends on several factors, primarily soil water content. Previous studies have shown that reductions in soil water content can lead to increases in canopy temperatures of up to 10 °C higher than air temperature (Luan and Vico 2021). Therefore, additional research is necessary to determine the critical threshold at which substituting T_air_ for T_c_ becomes inappropriate.

## 4. Conclusion

In this study, we investigated whether net radiation (R_n_) estimation could be improved by applying the Brunt approach for clear-sky emissivity, as previously calibrated by Formetta et al. (2016) and Li et al. (2017). Additionally, we examined whether air temperature (T_air_) could substitute canopy temperature (T_c_) in cotton crops under well-watered conditions. The results show that both clear-sky emissivity models improved R_n_ estimation compared with the original Allen equation, eliminating the need for site-specific calibration. Regarding the substitution of T_c_ with T_air_, we found that the improvement achieved by using T_c_ was modest, particularly at the daily time scale. Thus, under well-watered conditions, R_n_ can be satisfactorily estimated using T_air_ as a substitute for T_c_, especially at the daily time scale.

## Ethics

This work did not require ethical approval from a human subject or animal welfare committee.

## Data accessibility

Supplementary material and datasets in support of this work are available at https://zenodo.org/records/19353778

## Authors contributions

Thiago F. Duarte: Writing – original draft, Writing – review & editing, Conceptualization, Data curation, Visualization, Software, Methodology, Investigation, Formal analysis, Validation, Funding acquisition. Xuejun Dong: Writing – review & editing, Conceptualization, Methodology, Investigation, Data curation, Validation, Supervision, Resources, Funding acquisition, Project administration. Uzair Ahmad: Writing – review & editing, Data curation, Investigation. Noemi Tortorici: Writing – review & editing, Data curation, Investigation. Tonny J. A. Silva: Methodology, Validation, Writing – review & editing, Edna M. Bonfim-Silva: Methodology, Validation, Writing – review & editing, Daniel I. Leskovar: Methodology, Funding acquisition, Writing – review & editing.

## Acknowledgments

We appreciate Ray King, Bethany Speer, Sixto Silva, Dalton Thompson, Angela Jones, Joe Gonzalez and Randy Cox for assistance in crop management and field sample collection, and Christine Thompson and Liza Silva for administrative support. We thank the late Professor Dr. J. Tom Cothren for advice on cotton variety selection for the 2015 cotton trial and appreciate the help from Dale A. Mott for providing seeds for the 2025 cotton trial. We thank Dr. Charles Stichler for advice on sesame management.

