## Supplementary material for "Net radiation estimation using the Brunt equation for clear-sky emissivity and air and canopy temperatures for longwave radiation in well-watered crops"

**Appendix**

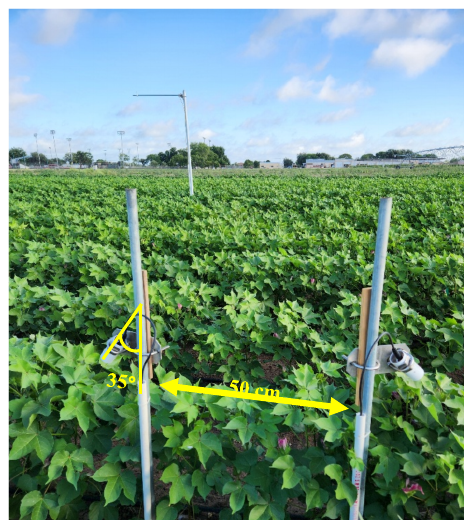

**Figure S1.** Details of the installation of the infrared radiometer sensors in the cotton field at the Uvalde Research Center in 2025. Shown in the background is the NR-Lite2 net radiometer mounted on a metal pole.

**Table S1.** The Brunt equation with different parameters calibrated by Formetta et al. (2016) and Li et al. (2017).

| Equation | Period | Reference | Local of calibration | Variable |
| --- | --- | --- | --- | --- |
| $\varepsilon_{c}=0.62 + 0.16\sqrt{e_{a}}$ | All day | Formetta | Texas, U.S. | e_a_ (kPa) |
| $\varepsilon_{c}=0.598 + 0.057\sqrt{e_{a}}$ | Daytime | Li | Seven stations Across U.S. | e_a_ (hPa) |
| $\varepsilon_{c}=0.633 + 0.057\sqrt{e_{a}}$ | Nighttime | Li |  | e_a_ (hPa) |

ɛ_c_: clear-sky emissivity; e_a_: water vapor pressure.
